# Does genetic risk help to predict amyloid burden in a non-demented population? A Bayesian approach

**DOI:** 10.1101/174995

**Authors:** Nicola Voyle, Willemijn Jansen, Aoife Keohane, Hamel Patel, Amos Folarin, Stephen Newhouse, Caroline Johnston, Kuang Lin, Pieter Jelle Visser, Angela Hodges, Richard JB Dobson, Steven J Kiddle, for the Alzheimer’s Disease Neuroimaging Initiative, EDAR and DESCRIPA study groups

**Affiliations:** MRC Social, Genetic and Developmental Psychiatry Centre, Institute of Psychiatry, Psychology and Neuroscience, King’s College London, London, UK; Department of Psychiatry and Neuropsychology, School for Mental Health and Neuroscience, Alzheimer Center Limburg, Maastricht University, The Netherlands; Department of Old Age Psychiatry, Institute of Psychiatry, Psychology and Neuroscience, King’s College London, London, UK; Department of Neurology and Alzheimer Center, VU University Medical Center, Amsterdam, The Netherlands; Helen Wills Neuroscience Institute, University of California, Berkeley; NIHR Biomedical Research Centre for Mental Health and Biomedical Research Unit for Dementia at South London and Maudsley NHS Foundation, London, UK; Farr Institute of Health Informatics Research, UCL Institute of Health Informatics, University College London, London, UK; MRC Biostatistics Unit, Cambridge Biomedical Campus, Cambridge Institute of Public Health, Forvie Site, Robinson Way, Cambridge, UK.

**Keywords:** Amyloid, Alzheimer’s Disease, Multi-modal, Polygenic Risk Score, Bayesian, Gene Expression Risk Score, Blood, Biomarker, Protein, Metabolite

## Abstract

**INTRODUCTION:** In this study we investigate the association between A*β* levels in cerebrospinal fluid (CSF) and genetic risk in a non-demented population. This paper presents the first analysis to use a Bayesian methodology in this area.

**METHODS:** Data from the Alzheimer’s Disease Neuroimaging Initiative (ADNI) and the EDAR* and DESCRIPA** studies was used in a Bayesian logistic regression analysis. We modeled CSF A*β* burden using age, diagnosis (healthy control or mild cognitive impairment), *APOE* and a polygenic risk score (PGRS) associated with Alzheimer’s Disease (AD). We compared models built using informative priors on age, diagnosis and *APOE* with non-informative priors on all variables.

**RESULTS:** The use of informative priors did not improve model performance in the majority of cases. Models using only age, diagnosis and *APOE* genotype showed the best predictive ability.

**DISCUSSION:** A previous study indicated that a PGRS of AD case/control status was associated with CSF A*β* burden in healthy controls. The current study suggests that this association does not lead to models that are more predictive of amyloid positivity than already known factors such as age and *APOE*.

*‘Beta amyloid oligomers in the early diagnosis of AD and as marker for treatment response’

**‘Development of screening guidelines and criteria for pre-dementia Alzheimers disease’

## 1 Introduction

It is hypothesized that late onset AD is caused by the presence of A*β* plaques in the brain tissue and hyperphosphorylated tau tangles in the neurons [1]. Hypothetical models and longitudinal studies indicate that A*β* pathology begins to develop up to 20 years prior to symptomatic changes [2, 3, 4]. This provides a window of opportunity for disease-modifying treatments, provided accurate and sensitive diagnostic tools are available. Existing tools for identifying the presence of pathology include measurements of amyloid and tau in CSF or in brain by the use of positron emission tomography (PET) imaging. Although these tests are reasonably accurate they are invasive and expensive. Peripheral biomarkers are sought as an intermediate step to provide cost-effective enrichment of people with high risk profiles for clinical trials, particularly secondary prevention trials of anti-amyloid therapeutics. In this study we aimed to address this by investigating the degree to which a genetic risk score was predictive of CSF A*β* in a non-demented population.

AD has been identified as a complex disease meaning its presence or absence is determined by environmental and genetic risk factors. Heritability is estimated to be 50-70% [5, 6, 7]. Over 20 risk loci have been identified for AD with Lambert *et al.* providing the most comprehensive Genome Wide Association Study (GWAS) to date [8, 9, 10, 11, 7]. However, it is estimated that currently identified SNPs can only explain 16-33% of phenotypic variation [9, 10, 7]. GWAS of A*β* endpoints are also highlighting promising results but sample sizes are considerably smaller so validation in larger numbers is needed [12]. An approach to consolidating the combined effect of several smaller genetic contributions to disease is found in polygenic risk scores (PGRS). A PGRS is calculated as a weighted sum and represents the cumulative effect of these smaller genetic effects. They have been shown to be informative in understanding genetic contributions to phenotypes beyond AD clinical diagnosis. For example, Sabuncu *et al.* found an AD case/control PGRS associated with A*β* burden in healthy controls (p < 0.0001). It has not been assessed whether this would provide predictive utility for enriching populations for prevention trials [13, 14, 15].

It is well known that model estimates, and often predictive ability, become more reliable as the number of individuals included in an analysis increases. However, in AD research the amalgamation of studies to create large datasets is often unfeasible due to differences in study populations and data collection methods. Initiatives such as the European Medical Information Framework (EMIF) are aiming to rectify this, but large populations with both multi-modal biomarker data and amyloid pathology measures are not yet available (www.emif.eu).

However, there is substantial information available on the associations between demographic variables and prevalence of A*β* burden, as discussed in meta-analyses by Jansen *et al.* and Ossenkoppele *et al.* [4, 16]. The former of these studies concentrated on persons without dementia (N=7583). The study concluded that age, *APOE* genotype and presence of cognitive impairment were associated with A*β* burden. No equivalent studies are available for tau pathology and consequently this work focuses on amyloid alone.

The present study uses data from the ADNI, EDAR and DESCRIPA cohorts, to investigate genetic risk as a blood biomarker of A*β* burden using a Bayesian methodology [17]. This study includes older individuals who do not have a clinical diagnosis of AD and are at a variable risk of developing the disease. This is the first study to use a Bayesian framework in AD blood biomarker research with the aim of investigating genetic risk as a marker in blood that could support strategies for identifying individuals at high risk of developing disease for recruitment into clinical trials. The models created combine age, diagnosis (control or MCI) and *APOE* genotype with a PGRS. A previous study has shown a case/control PGRS associated with A*β* in healthy controls. We aim to investigate whether this association translates to predictive ability. We hypothesized that by informing estimates for demographic variables using the large meta-analysis presented by Jansen *et al.* we would create more robust models. The Bayesian methodology used here has been made accessible to future researchers though the development of a simple graphical user interface (GUI).

## 2 Methods

### 2.1 Cohorts

#### 2.1.1 EDAR

EDAR is a prospective, longitudinal study with centers at multiple European sites. The study aims to examine and evaluate biomarkers of early AD and treatment response [17]. For more information see www.edarstudy.eu. Our access to samples and clinical and phenotypic information from the EDAR study was enabled by EMIF.

#### 2.1.2 DESCRIPA

DESCRIPA is a prospective, multi-center study based in Europe and coordinated by the European AD Consortium. The study focused on collecting data from non-demented subjects with the aim of developing screening guidelines and clinical criteria for AD in non-demented subjects. Further details of this study can be found in Visser *et al.* [18].

#### 2.1.3 ADNI

ADNI is a longitudinal cohort study aiming to validate the use of biomarkers in AD clinical trials and diagnosis. Data used in the preparation of this article were obtained from the ADNI database (adni.loni.usc.edu). The ADNI study was launched in 2003 as a public-private partnership, led by Principal Investigator Michael W. Weiner, MD. The primary goal of ADNI has been to test whether biological markers and clinical and neuropsychological assessment can be combined to measure the progression of mild cognitive impairment (MCI) and AD. For information, see www.adni-info.org. ADNI was approved by the institutional review boards of all participating institutions, and written informed consent was obtained for all participants.

The ADNI study comprises three stages. ADNI 1 is the initial study (target N = 800). ADNI GO contains a subset of the controls and MCI participants from ADNI 1 and is supplemented by additional individuals with MCI (target N = 700). ADNI 2 enhances ADNI 1 and ADNI GO further with the inclusion of new participants in all diagnostic groups (target N = 1350).

This study uses data from ADNI 1 and the ADNI 2 and ADNI GO sub-groups, referred to as ADNI 2 from here onwards.

### 2.2 Genetics

Samples from EDAR and DESCRIPA were genotyped on the Illumina HumanOmniExpressExome-8v1.2 BeadChip and processed together (N = 336) [19]. This BeadChip contains 960,919 markers of which 273,000 represent functional exomic markers. The data was processed in Genome Studio (as described at bit.ly/1VpRclH) before being run through the rare variant caller Zcall [20] (as described here: bit.ly/1YKHYhK). ADNI 1 samples were run on the Human610-Quad BeadChip (N = 818) while ADNI 2 and ADNI GO samples were run on the Illumina HumanOmniExpress BeadChip (N = 432). The HumanOmniExpress BeadChip is similar to that used in the EDAR and DESCRIPA studies but does not include exomic markers, while the Human610-Quad BeadChip is older. Details of the genotyping protocols followed in ADNI are given elsewhere [21].

The cohorts were subject to quality control and imputation separately, as described in Coleman *et al.* [22]. In short, the data was filtered to ensure a minor allele frequency of greater than 5% for all SNPs before removal of rare variants and subjects with high levels of missing data. SNPs that differed significantly (p < 0.00001) from the Hardy-Weinberg equilibrium were removed. The data was pruned for SNPs in linkage disequilibrium and for genetically similar individuals. Finally, the data was imputed using reference files from the 1000 Genomes Project. [23]

### 2.3 Amyloid measurements

Throughout this study amyloid measurements in cerebrospinal fluid (CSF) are used as the endpoint of interest. All A*β* measurements were dichotomized as detailed in the meta-analysis from Jansen *et al.*. The distribution of amyloid burden in all studies was bimodal (as expected) making this dichotomization biologically relevant. Low CSF A*β* is referred to as ‘abnormal’ A*β* burden while high CSF A*β* is referred to as ‘normal’. The details for each study are as follows:

#### 2.3.1 EDAR

Details of CSF collection and analysis can be found at www.edarstudy.eu. In brief, CSF measurements were collected using the Alzbio3 Luminex assay in one batch at the end of the study. CSF amyloid measurements were dichotomized at the previously published threshold of 389pg/ml.

#### 2.3.2 DESCRIPA

Details of CSF measurements in DESCRIPA have been described elsewhere [19]. In brief, CSF measurements were analyzed in one laboratory and collected using single-parameter ELISA kits (Innogenetics, Ghent, Belgium). CSF amyloid samples were dichotomized using the previously published threshold of 550pg/ml.

#### 2.3.3 ADNI

For ADNI, datasets used to extract CSF measures of amyloid were chosen to maximize sample size. The dataset ‘UPENNBIOMK2’ was used for ADNI 1 and ‘UPENNBIOMK6’ for ADNI 2 and ADNI GO. Both datasets contain CSF measurements collected using the xMAP Luminex platform and Innogenetics immunoassay kits. The CSF measures were dichotomized at the previously published threshold (192 pg/ml).

### 2.4 Statistical analysis

All statistical analysis was performed in R version 3.1.1 [24]. Models were built in ADNI 1 data and tested in data from EDAR, DESCRIPA and ADNI 2. We built models including age, diagnosis (healthy control or MCI) and *APOE* genotype as covariates (‘basic model’) and models including these variables with the addition of a PGRS (‘PGRS model’). The predictive ability of each model was quantified using accuracy, sensitivity, specificity and area under the Receiver Operating Characteristic (ROC) curve [25, 26, 27].

#### 2.4.1 PGRS

PGRS were created using the software package PRSice [28]. Effect sizes from stage 1 of the International Genomics of Alzheimer’s Project (IGAP) case/control GWAS were used as the weights to generate the risk score (N = 54,162, number of SNPs = 7,055,881) [8]. We used 0.5 as the p-value threshold for inclusion in the PGRS. This threshold showed the most significant association with case/control diagnosis in the large IGAP PGRS study [15]. The genetic region coding for *APOE* was removed from all scores and included as a covariate in modeling.

#### 2.4.2 Data analysis

This study aimed to predict dichotomized amyloid burden using genetic risk in a Bayesian logistic regression. The method was implemented using the R function ‘MCMClogit’ in the ‘MCMCpack’ package [29]. Models were built using a Metropolis sampler with 100,000 MCMC iterations, with the first 3,000 discarded as burn-in. This number of iterations ensured the ratio of standard deviation to Monte Carlo Standard Error (MCSE) was less than 5% for all parameters.

The variables included in the ‘basic model’ were chosen based on the meta-analysis published by Jansen *et al.* [4]. The study used generalized estimating equations (GEEs) to predict amyloid burden from age, diagnosis (control or MCI) and *APOE ε*4 status (defined as the presence or absence of any number of *E*4 alleles). Age is centered at 70 years. In this study we only consider healthy controls and people with a diagnosis of MCI. The best model identified by Jansen *et al.* was:

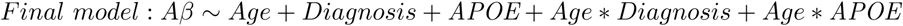

We have used the estimates from this meta-analysis to inform the regression estimates of variables in this study where possible. It can be shown that the estimates from GEEs are normally distributed [30] and hence we included these estimates as priors using a multivariate normal distribution (see Table 1). The PGRS had a non-informative multivariate normal prior with mean 0 and variance 100. We also created models where all variables had the non-informative Normal(0,100) priors for comparison.

**Table 1:** Informative prior distributions

| Variable | Estimate | SE |
| --- | --- | --- |
| Intercept | 0.879 | 0.0967 |
| Age | 0.064 | 0.006 |
| Diagnosis = Control | -0.964 | 0.0793 |
| *APOE* status = 0 | -1.493 | 0.0772 |
| Age * *APOE* status = 0 | -0.021 | 0.0079 |
| Age * Diagnosis = Control | 0.019 | 0.0081 |

SE = Standard error.
MCI is used as the reference diagnosis.
*APOE* status 1 (at least one $\epsilon 4$ allele) is used as the reference level.

The tuning parameter of each model was adjusted to achieve an acceptance rate in the Metropolis sampler of approximately 0.35. This tuning was performed over the 3,000 burn-in samples. This acceptance rate is slightly higher than advised in the literature as lower acceptance rates were causing reduced mixing [31].

#### 2.4.3 Graphical User Interface

The analysis methods used in this study have been packaged into a user-friendly application through Rshiny [32]. The application is available to download from https://github.com/KHP-Informatics/bayesian-logistic-regression-r-shiny-app.git.

## 3 Results

### 3.1 PGRS

In the ADNI cohort the genetic data was imputed from 479,073 and 599,526 SNPs in ADNI 1 and ADNI 2 respectively, to 8,799,802 and 6,336,499 SNPs. In EDAR and DESCRIPA the data was increased from 619,609 SNPs to 5,409,779 by imputation. The PGRS was standardized by subtracting the mean and dividing by the standard deviation, per cohort.

### 3.2 Cohort demographics

The demographics given below are from the individuals in ADNI 1, ADNI 2, EDAR and DESCRIPA with CSF and GWAS data available.

In ADNI 1 (training data) there is 1 point difference in median MMSE between subjects with normal and abnormal A*β* (28 vs. 29). However, the larger sample size of ADNI 1 (compared to ADNI 2, EDAR and DESCRIPA) means this difference is statistically significant. Diagnosis and *APOE* genotype also show significant differences between groups, as we would expect. We see a significant difference (p-value < 0.05) in the PGRS between groups. Similar associations are seen in the data from EDAR and DESCRIPA with age also being nominally significantly associated with normal and abnormal A*β*. ADNI 2 data shows no significant difference in any demographic variable. This is likely to be due to the small sample size (N=43) of this population.

### 3.3 Data analysis

The test data indicated that, in most cases, the addition of informative priors on age, diagnosis and *APOE* genotype did not improve the predictive ability of models. Furthermore, no model including a PGRS showed higher accuracy than the basic models. The basic model with non-informative priors achieved the highest accuracies at 54% and 49% with a high sensitivity of 81% and 63%, but low specificity (23% and 41%). See Table 3 for full results.

**Table 2:** Cohort demographics

| Demographic | ADNI 1 (N=272) |  |  | ADNI 2 (N=43) |  |  | EDAR and DESCRIPA * (N=127) |  |  |
| --- | --- | --- | --- | --- | --- | --- | --- | --- | --- |
| | Normal CSF Aβ | Abnormal CSF Aβ | P-value | Normal CSF Aβ | Abnormal CSF Aβ | P-value | Normal CSF Aβ | Abnormal CSF Aβ | P-value |
|  | (N = 105) | (N = 167) |  | (N = 27) | (N = 16) |  | (N = 60) | (N = 67) |  |
| Median age [IQR] | 74.1 [8.10] | 75.1 [8.05] | 0.837 | 67.8 [12.55] | 73.75 [11.075] | 0.087 | 66 [8.75] | 69.55 [11.83] | 0.047 |
| Gender (%): |  |  | 0.61 |  |  | 0.528 |  |  | 0.859 |
| Female | 36.2 | 39.5 |  | 51.9 | 37.5 |  | 46.7 | 49.3 |  |
| Male | 63.8 | 60.5 |  | 48.1 | 62.5 |  | 53.3 | 50.7 |  |
| Median years in education [IQR] | 16 [4] | 16 [4] | 0.955 | 16 [4] | 16[4] | 0.949 | 12 [7] | 10 [5] | 0.334 |
| Median MMSE [IQR] | 29 [3] | 28 [3] | <0.001 | 29 [2] | 27.5 [2.25] | 0.1 | 28.5 [3] | 27 [5] | 0.024 |
| Diagnosis (%) |  |  | <0.001 |  |  | 0.446 |  |  | 0.002 |
| MCI | 42.9 | 76.6 |  | 74.1 | 87.5 |  | 71.7 | 92.5 |  |
| CTL | 57.1 | 23.4 |  | 25.9 | 12.5 |  | 28.3 | 7.5 |  |
| *APOE* status (%) |  |  | <0.001 |  |  | 0.343 |  |  | 0.004 |
| 0 | 84.8 | 38.3 |  | 66.7 | 50.0 |  | 66.7 | 40.3 |  |
| 1 | 15.2 | 61.7 |  | 33.3 | 50.0 |  | 33.3 | 59.7 |  |
| Median PGRS [IQR] | -0.615 [2.196] | -0.560 [2.095] | 0.018 | -0.088 [1.362] | -0.221 [1.127] | 0.763 | -0.252 [1.387] | 0.226 [1.266] | 0.022 |

Kruskal Wallis chi-squared was used to test between normal and abnormal groups for continuous demographic variables.
Fishers exact was used to test between normal and abnormal groups for categorical demographic variables.
*APOE* status is 1 if an individuals genotype contains any $\epsilon 4$ alleles, and 0 otherwise.
IQR = Inter-quartile range; CSF = Cerebrospinal Fluid; MMSE = Mini Mental State Exam; MCI = Mild Cognitive Impairment; CTL = Control.
\* One individual has missing MMSE and education information.

**Table 3:** Test data results

| Model | Informative priors? | Accuracy [95% CI] | Sensitivity | Specificity | AUC ROC |
| --- | --- | --- | --- | --- | --- |
| **EDAR and DESCRIPA** |  |  |  |  |  |
| Demographics | No | 0.535 [0.445; 0.624] | 0.806 | 0.233 | 0.412 |
| Demographics | Yes | 0.543 [0.453; 0.632] | 0.776 | 0.283 | 0.412 |
| PGRS | No | 0.457 [0.368; 0.547] | 0.657 | 0.233 | 0.391 |
| PGRS | Yes | 0.394 [0.308; 0.484] | 0.493 | 0.283 | 0.394 |
| **ADNI 2** |  |  |  |  |  |
| Demographics | No | 0.488 [0.333; 0.645] | 0.625 | 0.407 | 0.458 |
| Demographics | Yes | 0.488 [0.333; 0.645] | 0.625 | 0.407 | 0.461 |
| PGRS | No | 0.465 [0.312; 0.623] | 0.625 | 0.37 | 0.396 |
| PGRS | Yes | 0.442 [0.291; 0.601] | 0.5 | 0.407 | 0.4 |

CI = Confidence Interval; AUC ROC = Area under the Receiver Operating Characteristic Curve;
PGRS = Polygenic Risk Score

## 4 Discussion

This study shows that the predictive ability of models including age, diagnosis and *APOE* genotype is not improved by the addition of a PGRS despite previous studies showing an association [13]. The PGRS used in this analysis was trained on a case/control endpoint. As GWA studies of amyloid endpoints become available the predictive ability of a PGRS trained using this information is likely to improve [33].

This paper presents the first analysis to use a Bayesian methodology in AD blood biomarker research. We aimed to inform estimates of well-researched risk variables (age, *APOE* status and control or MCI status) by including prior information from a large meta-analysis [4]. In this study we see that this approach does not improve predictive ability of models over those without informative prior information. However, we have only used one type of Bayesian methodology. There is a risk that if ill-fitting priors are used in combination with small sample sizes, model fit may be driven by the prior distributions. It is possible that the priors used here are not optimal as our population demographics are slightly different from those seen by Jansen *et al.* [4]. However, we believe informing model estimates with information from previous literature is likely to reduce false positives in biomarker studies of a small size. Bayesian analysis is one way of doing that; other methods should also be investigated.

The creation of the PGRS also created some limitations for this study. Firstly, the PGRS used here only utilizes common genetic variation. This is because rare genetic variants, such as *TREM2*, may not be significantly associated with disease in small populations. As larger studies become available inclusion of such variants should be investigated. Further, the PGRS was created using a simple additive method. This simplistic method is likely to be sub-optimal and as new methodologies for creating PGRS become available they should be investigated in this setting. Additionally, the platforms used to measure genetics differed between the discovery and test cohorts used in this study. It is well documented that there can be inconsistencies between *omics* platforms which could have contributed to the reduced predictive ability seen here [34, 35, 36, 37].

It is important to bear in mind in all further research that differences in normal/abnormal CSF A*β* cut-offs can make replication and research in other cohorts difficult. Work is being done to investigate these values with an aim of standardization [38]. This study also has the limitation that the IGAP data (used to generate the PGRS) is not independent of the training and test data, as ADNI was included in IGAP. However, we believe the benefit of larger sample size outweighs this. Finally, the cohorts used in this study are still of a relatively small size. Although we tried to address this through the use of Bayesian methods, studies of larger sample sizes will be vital for further investigations of blood based biomarkers of A*β*.

This study also presents opportunities for further work. Firstly, it is possible that markers identified in case/control studies are associated with other AD related phenotypes such as tau burden. If large meta-analyses of the risk factors affecting tau burden, and other endophenotypes, become available it would be interesting to perform similar Bayesian analysis on these alternative endpoints. Secondly, in the setting of preventative clinical trials the most common way to measure brain amyloid burden is through the use of a PET scan. In this study we have used CSF. However, it has been shown that measurements from CSF and PET are highly correlated meaning markers identified for one can reasonably be tested to see whether they are transferable to the other. In this study the use of CSF allowed us to maximize the sample size with measurements of amyloid and genetics. If promising blood markers of CSF measures are identified, future studies should perform similar analysis using a measurement of A*β* derived from a PET scan. Furthermore, the effect of a measure of brain reserve on model accuracy could be investigated. It is well-known that some people with high levels of brain amyloid burden at autopsy show no cognitive deficit during their lifetime. It has been shown in recent studies that levels of ‘brain reserve’ may be driving the difference between people who have high levels of pathology and no symptoms and those who show symptoms [39]. This is motivating a theory that increased brain reserve may prevent development of symptoms even if pathology is present. Although brain reserve itself may be hard to quantify it may be possible to measure and model associated lifestyle, environmental and psychological factors such as social networks.

Although there are further candidates and alternative methods to be considered in the search for a blood biomarker of amyloid burden it is imperative that appropriate populations are used. As in this work, the search in asymptomatic individuals is likely to have the biggest impact for enrichment of clinical trials. Furthermore, rigorous testing of biomarkers is essential. Without replication it is probable that model performance is overestimated.

## 5 Conclusions

This paper presents the first analysis to use a Bayesian methodology in the search for a blood biomarker of AD. We see that the Bayesian approach does not improve predictive ability in this setting, and that *omics* measurements do not improve the predictive ability of models above that of demographics alone. We have been unable to demonstrate any additional benefit over age diagnosis and *APOE* genotype by including a case/control PGRS when predicting amyloid positivity in subjects without a clinical diagnosis of AD or other dementia.

## 6 Acknowledgements

This work represents independent research part funded by the National Institute for Health Research (NIHR) Biomedical Research Centre at South London and Maudsley NHS Foundation Trust and King’s College London. This study was supported by researchers at the National Institute for Health Research University College London Hospitals Biomedical Research Centre. The research leading to these results has received support from the Innovative Medicines Initiative Joint Undertaking under grant agreement number 115372, resources of which are composed of financial contribution from the European Union’s Seventh Framework Programme (FP7/2007-2013) and EFPIA companies’ in kind contribution. Steven Kiddle is supported by an MRC Career Development Award in Biostatistics (MR/L011859/1). Nicola Voyle is funded by the Alzheimer’s Society. The views expressed are those of the author(s) and not necessarily those of the NHS, the NIHR or the Department of Health.

The authors would like to thank Dr. Steven Hill and Dr. Robert Goudie from the MRC Biostatistics Unit for their guidance in Bayesian methods.

This work was also supported by awards to establish the Farr Institute of Health Informatics Research at UCL partners, London, from the Medical Research Council, Arthritis Research UK, British Heart Foundation, Cancer Research UK, Chief Scientist Office, Economic and Social Research Council, Engineering and Physical Sciences Research Council, National Institute for Health Research, National Institute for Social Care and Health Research, and Wellcome Trust (grant MR/K006584/1).

Data collection and sharing for this project was funded by the Alzheimer’s Disease Neuroimaging Initiative (ADNI) (National Institutes of Health Grant U01 AG024904) and DOD ADNI (Department of Defense award number W81XWH-12-2-0012). ADNI is funded by the National Institute on Aging, the National Institute of Biomedical Imaging and Bioengineering, and through generous contributions from the following: AbbVie, Alzheimers Association; Alzheimers Drug Discovery Foundation; Araclon Biotech; BioClinica, Inc.; Biogen; Bristol-Myers Squibb Company; CereSpir, Inc.; Eisai Inc.; Elan Pharmaceuticals, Inc.; Eli Lilly and Company; EuroImmun; F. Hoffmann-La Roche Ltd and its affiliated company Genentech, Inc.; Fujirebio; GE Healthcare; IXICO Ltd.; Janssen Alzheimer Immunotherapy Research and Development, LLC.; Johnson and Johnson Pharmaceutical Research and Development LLC.; Lumosity; Lundbeck; Merck and Co., Inc.; Meso Scale Diagnostics, LLC.; NeuroRx Research; Neurotrack Technologies; Novartis Pharmaceuticals Corporation; Pfizer Inc.; Piramal Imaging; Servier; Takeda Pharmaceutical Company; and Transition Therapeutics. The Canadian Institutes of Health Research is providing funds to support ADNI clinical sites in Canada. Private sector contributions are facilitated by the Foundation for the National Institutes of Health (www.fnih.org). The grantee organization is the Northern California Institute for Research and Education, and the study is coordinated by the Alzheimer’s Disease Cooperative Study at the University of California, San Diego. ADNI data are disseminated by the Laboratory for Neuro Imaging at the University of Southern California.

